# Allogenic platelet-rich plasma and platelet-rich plasma extracellular vesicles alter the proteome of tenocytes in an in vitro equine model of tendon inflammation: A pilot study

**DOI:** 10.1101/2024.08.27.609926

**Authors:** Emily J Clarke, Anders Jensen, Alexandra M Gillen, David Bardell, Mark Senior, James R Anderson, Mandy J Peffers

## Abstract

**Background:** Tendon injuries are common in horses, often resulting in a high risk of reinjury. Hemoderivative therapeutics including platelet-rich plasma (PRP) show promise but outcomes vary due to inconsistent composition. PRP is rich in soluble growth factors and extracellular vesicles (EVs), the latter facilitate cell-to-cell communication by delivering biologically active cargo. This pilot study profiled the proteome of PRP and PRP-derived EVs and examined their effects on an equine tendon inflammatory model *in vitro*.

**Methods:** Plasma was isolated via double centrifugation and PRP produced using a commercial filtration kit. EVs were isolated from PRP and plasma using differential ultracentrifugation and characterised with the Exoview Tetraspanin assay. Equine tenocytes were stimulated with interleukin 1β and tumor necrosis factor α, then treated with PRP or PRP-derived EVs. Proteomic analysis was conducted on cell lysates, PRP and PRP-EVs using data-dependent acquisition liquid chromatography-tandem mass spectrometry and the data analyzed using multivariate and univariate approaches.

**Results:** PRP contained 575 quantifiable proteins and PRP-derived EVs 209 proteins. When compared to plasma and plasma derived EVs respectively, PRP and PRP-EVs were enriched in proteins associated with cellular waste disposal and inhibition of lipid metabolism. Experimental factors (inflammatory stimulation and/or treatments) significantly affected the abundance of 18 proteins as expressed in equine tenocytes including col1a1 (col1a1) and sequestosome 1, associated with collagen metabolism and nuclear factor kappa B signaling.

**Discussion:** The findings from this study suggest PRP-derived EVs influence inflammatory tenocytes and may be crucial to the efficacy of PRP.

## 1. Introduction

Equine musculoskeletal injuries are highly prevalent within the performance horse population, accounting for 82% of all injuries to racehorses competing in National Hunt and flat races. Of these, 46% involved tendon or ligaments ^1,2^, with the superficial digital flexor tendon (SDFT), an energy storing tendon, most prone. The SDFT experiences large forces and repetitive loads ^3^ accumulating microdamage, due to exercise and ageing, predisposing it to rupture ^4^. Risk factors for tendon injury include breed, sex, age, weight and variations in tendon vascular supply ^5–8^. A significant issue with SDFT damage is the incidence of reinjury ^9^ and reduced biomechanical fortitude ^10^ which results in compromised welfare, pain and lameness.

Tendon is composed of a fibrous extracellular matrix (ECM) accompanied by a small number of resident tenocytes that maintain structure through the production of matrix molecules^1^. The structure is largely avascular, postulated to promote poor healing ^11^. Significant inflammation often accompanies injury, with many pro-inflammatory cytokines expressed including tumour necrosis factor alpha (TNF-α) and interleukin 1 beta (IL-1β) ^12,13^. Tendon healing is often slow due to the poor innate regenerative properties. The repaired tissue is characterised by fibrosis and collagen type III, producing an inferior structure; contributing towards a high re-injury rate ^14^.

Heamoderivative therapeutics, including autologous platelet-rich plasma (PRP), have become a popular form of treatment for orthopaedic conditions including tendon injury. This is due to cost, ease of preparation, minimal equipment requirement and its immunomodulatory effect ^15^. Additionally, it promotes haemostasis, anti-inflammatory cytokine release and the release of growth factors such as platelet-derived growth factor (PGF), transforming growth factor-β1 (TGFβ1), transforming growth factor-β2 (TGFβ2), vascular endothelial growth factor (VEGF), basic fibroblastic growth factor (FGF), and epidermal growth factor (EGF), following degranulation of alpha platelet granules ^15–17^. PRP is defined as having at least twice the platelet count of baseline whole blood values ^18,19^. In a randomised prospective trial using PRP a single intralesional treatment up to 8 weeks after the onset of clinical signs of tendon injury reduced lameness and advanced the organisation of repair tissue ^20^. Bosch et al demonstrated PRP had the capacity to increase metabolic activity and improve the maturation of repaired tissue in experimentally induced tendon lesions ^21^. However, PRP does present some drawbacks as clinical variables effect the autologous product composition and level of activation ^21^. In addition, the lack of standardised PRP composition has contributed to variable clinical outcomes and a lack of reproducibility ^20,22^.

Extracellular vesicles (EVs) have become an area of interest in understanding the mechanistic action of PRP ^23^. EVs are nanoparticles secreted by cells, enveloped in a phospholipid bilayer and are involved in intercellular communication by transporting biologically active cargo between cells ^24^. They are categorised by size, density and biochemical composition. Graca *et al* ^25^ explored the effect of platelet-derived EVs in a bioengineered model of human tendon injury and observed an increase in tenogenic markers, subsequently promoting a healthy ECM phenotype, remodeling and increasing the synthesis of anti-inflammatory mediators. Thus, EVs derived from platelets have the capacity to transfer platelet-derived content to cellular recipients and organs inaccessible to platelets ^26,27^. Furthermore, EVs have a low immunogenicity and are easily stored and obtained. Pre-clinical studies are aiming to quantify the effect of PRP-EVs therapeutically to determine if they are a viable superior alternative to cell-based regenerative therapeutics ^28^. Therefore, it is important to understand the composition of PRP and PRP-EVs and the effect these EVs have in mediating the therapeutic effect observed, to develop a restorative product that can deliver improved and reproducible clinical outcomes as a blood-based product or EVs alone. Our pilot study hypothesised that PRP and PRP-EV treatment of cytokine-stimulated tenocytes would result in a change in the cellular phenotype at the protein level.

## 2. Materials and Methods

### 2.1 Sample Collection

SDFT tendons were collected from Thoroughbreds (n=5) and an Irish Sports horse (n=1) under ethical approval and owner consent (VREC561). Four mares and two geldings were included with a mean age of 10.7 (standard deviation, SD ±4.07) years old.

### 2.2 Whole Blood Collection

Venous whole blood was collected post-mortem from an abattoir (n=3) into acid citrate dextrose tubes (ACD-A). Samples were inverted and kept at room temperature until PRP isolation. Samples were collected from an abattoir as a by-product of the agricultural industry and processed within 12 hours of euthanasia. The Animals (Scientific Procedures) Act 1986, Schedule 2, does not define collection from these sources as a scientific procedure.

### 2.3 PRP Isolation and Characterisation

PRP was isolated using the V-PET Equine Platelet Enhancement Therapy System (Vet Stem Biopharma, Poway, California) following the manufacturer’s guidelines. 55 ml of whole venous blood was used to produce 8 ml of PRP. Whole blood and PRP from each of the three donors were analysed using a IDEXX ProCyte Dx haematology analyser (IDEXX, Hoofddorp, the Netherlands) to obtain a haematology profile and ensure PRP has >2x platelet count versus baseline whole blood values ^18,19^.

### 2.4 EV Isolation

Equine PRP samples (100 μl) underwent differential ultracentrifugation (dUC) to isolate EVs, as previously described ^29,30^. All isolation and characterisation procedures were uploaded to the EV-TRACK database; EV-TRACK ID EV230986.

### 2.5 EV Characterisation – Exoview

The ExoView platform (NanoView Biosciences, Malvern Hills Science Park, Malvern) was used to determine EV concentration and surface marker identification (CD9, CD81 and CD63). Samples were prepared and analysed as previously described ^29,30^.

### 2.6 Cell Isolation and Experimental Design

SDFT explants were dissected from the forelimbs of donors and expanded as previously described ^31^. Equine tenocytes (P3) were grown in 12 well plates in phenol red-free complete media, with the addition of L-glutamine, until 70% confluent. Nine plates were used in total, six for technical duplicates for qPCR analysis and three for proteomic analysis. Upon 70% confluency, cells were washed in PBS and serum starved for 24 hours. Following this, tenocytes were treated with cytokines; 10 ng/ml of IL-1β and TNF-α (Recombinant Equine IL-1β, R&D Systems, Abingdon, UK) for 24 hours. After treatment of cytokines to induce a model of tendon inflammation cells received no treatment (control) or were treated with PRP (100 µl) and PRP-EVs (EVs derived from 100 µl of PRP, (∼500,000 EVs per µl), (3.3%(v/v%)) accordingly and incubated for an additional 24 hours at 37°C in hypoxic conditions. The experimental groups were: (1) control, (2) control + PRP, (3) control +PRP EVs, (4) cytokine stimulation, (5) cytokine stimulation + PRP and (6) cytokine stimulation + PRP EVs. Nine plates were used in total, six for technical duplicates for qPCR analysis and three for proteomic analysis.

For RNA extraction analysis, tenocytes were washed with PBS, suspended in Trizol (Invitrogen, Paisley, UK), scraped and stored at -80°C for later RNA extraction. Cells for proteomic analysis were washed, trypsinised as previously described ^31^, counted, centrifuged at 2400 rpm and stored in 25 mM ammonium bicarbonate (AmBic) (Sigma-Aldrich, Dorset, UK) at -80°C. A summary of the experimental design is shown in Figure 1.

**Figure 1.**
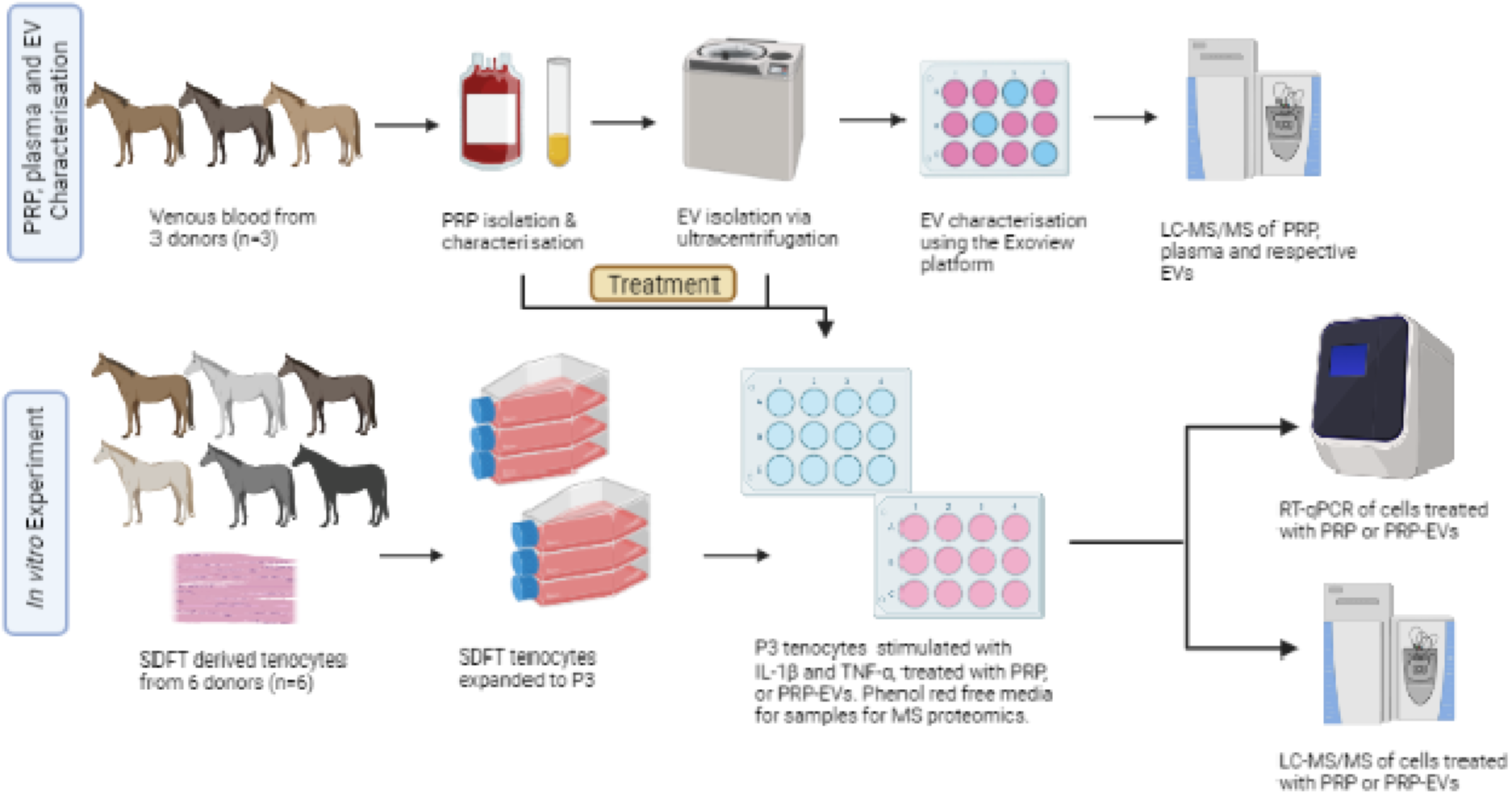
Experimental design overview. Equine venous blood (n=3) was used to produce PRP. PRP was characterised, PRP-EVs isolated using differential ultracentrifugation and the proteome quantified. The profiled PRP and PRP-EVs were then used in an *in vitro* equine tendon inflammatory model following cytokine stimulation to ascertain the cellular response to PRP and PRP-EVs. This illustration was created in Biorender.com.

### 2.8 RNA Extraction

RNA was extracted following a phase extraction protocol ^32,33^. Samples were suspended in 500 µl of Trizol and RNA extracted as previously described ^34^. Total RNA was measured using a spectrometer (NanoDrop™ 1000 Spectrophotometer, Thermo Fisher Scientific, Waltham, US) and samples stored at -80°C.

### 2.9 cDNA Synthesis and Reverse Transcription-quantitative Polymerase Chain Reaction

For cDNA synthesis all reagents were from Promega, Southampton, UK, unless otherwise stated. Total RNA (500 ng) was suspended in RNase-free water for cDNA synthesis. 9µl was added to 1µl random primer and pulse centrifuged. Samples were placed in a thermocycler (Applied biosystems Veriti, 96 well thermocycler, Warrington, United Kingdom) and incubated at 70°C for 5 minutes, then 4°C for 2 minutes. Each sample had Moloney murine leukemia virus reverse transcriptase (M-MLV RT, Promega), M-MLV RT 5x reaction buffer and dNTP (Bioline Reagents Ltd., London, UK) added was incubated at 37°C for 1 hour then 95°C for 10 minutes.

### 2.10 Reverse Transcription-quantitative Polymerase Chain Reaction

cDNA was diluted to 5 ng/µl and 5 µl of sample was pipette per well into a 96 well PCR plate in technical duplicates. Each sample was added to a Master mix comprised of Takyon No ROX SYBR 2x master mix blue dTTP (Eurogentec, Liege, Belgium), and a 100mM concentration of appropriate forward and reverse primers (Eurogentec, Liege, Belgium). qPCR analysis was performed on a Roche Lightcycler 480, samples were preincubated at 95°C for 3 minutes, followed by amplification for 40 cycles; denaturation at 95°C for 10 seconds then annealing at 60°C for 1 minute. At the end of cycling a melt curve was produced. All exon spanning primers used were designed using the NCBI PrimerBlast data base and were species specific. Relative quantification was used in order to process raw cq values in order to produce relative gene expression values using 2-Δct ^35^. Glyceraldehyde 3-phosphate dehydrogenase (GAPDH (F: GCATCGTGGAGGGACTCA, R: GCCACATCTTCCCAGAGG)) was selected as a housekeeping gene ^36^, collagen type I alpha 2 chain (COL1A2 (F: GCACATGCCGTGACTTGAGA, R: CATCCATAGTGCATCCTTGATTAGG)), insulin like growth factor binding protein 6 (IGFBP6 (F: GAACCGCAGAGACCAACAGA, R: ACGGGCCCATCTCCGT)) and matrix metalloproteinase (MMP3 (F: TCTTGCCGGTCAGCTTCATATAT, R: CCTATGGAAGGTGACTCCATGTG) and MMP13 (F: GAGCATCCTCCCAAAGACCTT, R: CATAACCATTAAGAGCCCAAAATT)).

### 2.11 Protein Extraction

For cellular lysates the supernatant was removed from cells and the cellular pellet dissolved in 100 µl of 25 mM AmBic (Sigma Aldrick, Dorset, UK) as previously described^37,38^.

PRP and plasma EVs pellets were suspended in 200μl of urea lysis buffer (6M Urea (Sigma-Aldrich, Dorset, United Kingdom), 1 M AmBic (Fluka Chemicals Ltd., Gillingham, UK) and 0.5% sodium deoxycholate (Sigma-Aldrich, Dorset, United Kingdom)). Protein was extracted as previously described ^39,40^ and quantified with a Pierce 660 nm assay following the manufacturer’s guidelines ^41^.

### 2.13 Protein Equalisation Using Proteominer™ Columns

Proteominer™ columns were used to reduce the protein concentration dynamic range for PRP and plasma samples as previously described ^42–44^. Sample protein was subsequently bound to beads and used for on bead protein digestion ^45^.

### 2.14 Protein Digestion of PRP-EVs, Plasma-EVs and Cell Lysates

PRP-EVs, plasma-EVs and tenocyte cell lysates were suspended in 160 μl of 25mM AmBic and were trypsin digested in-solution as previously described ^39,40^.

### 2.15 Label Free Liquid Chromatography Tandem Mass Spectrometry

Data-dependent liquid chromatography tandem mass spectrometry (LC-MS/MS) analyses were conducted on a QExactive HF Quadrupole-Orbitrap mass spectrometer (Thermo Scientific, Hemel Hempstead, UK) coupled to a Dionex Ultimate 3000 RSLC nano-liquid chromatograph (Thermo Scientific, Oxford, United Kingdom) ^39,40,42,44,46,47^ .

For label-free quantification, raw files of the acquired spectra were analysed using ProgenesisQI™ software (Waters, Manchester, UK). A local Mascot server (Version 2.8.2) was used to identify peptides, searching against the Unihorse database with carbamidomethyl cysteine as a fixed modification and methionine oxidation as a variable modification, peptide mass tolerance of 10 ppm and a fragment mass tolerance of 0.01 Da. Data were deposited to the ProteomeXchange Consortium via the PRIDE^48^ (PXD046066 and 10.6019/PXD046066).

### 2.16 Statistical Analysis

Statistical analyses were undertaken in GraphPad Prism 10.0 (California, USA), R and Metaboanalyst V5.0 ^49^. Normality tests were performed following Shapiro-Wilk analysis where applicable followed by appropriately selected univariate and multivariate statistical tests and corrected for multiple testing using post hoc Tukey tests. Statistical significance was attributed to a p<0.05. The proteomics data associated with PRP, plasma and associated EVs were quality controlled; proteins with more than 50% missing observations were removed from the dataset. Proteomics data associated with the *in vitro* experiment were also quality controlled; proteins with 25% or more missing observations were removed from the dataset, as were contaminating blood-associated proteins. The remaining data was normalised using probabilistic quotient normalisation (PQN) of the control group and log-transformed (base 10) for downstream analysis.

### 2.18 Functional Enrichment Analysis

Functional enrichment analysis was performed using Ingenuity Pathway Analysis (IPA; Qiagen, Hilden, The Netherlands) as previously described with all proteins identified used as background ^40^.

## 3. Results

### 3.1 PRP-EVs were enriched in CD9

PRP-EVs from a pooled sample of the previously analysed PRP (data shown in supplementary figure 1) were characterised using the Human Exoview Tetraspanin Assay. PRP-EVs expressed CD81 and CD9 (Figure 2A and 2B). CD63 was not reported due to poor protein homology between equine and human CD63 tetraspanin 50. PRP-EVs had a concentration of 5.23×10^10^ particles/ml and plasma EVs had a concentration of 6.91×10^10^ particles/ml. As plasma EVs were not used in the *in vitro* equine tendon inflammatory model experiment, their characterisation data is not shown. PRP-EVs were visualized with fluorescent microscopy, highlighting tetraspanin expression and EV morphology (Figure 2C).

**Figure 2.**
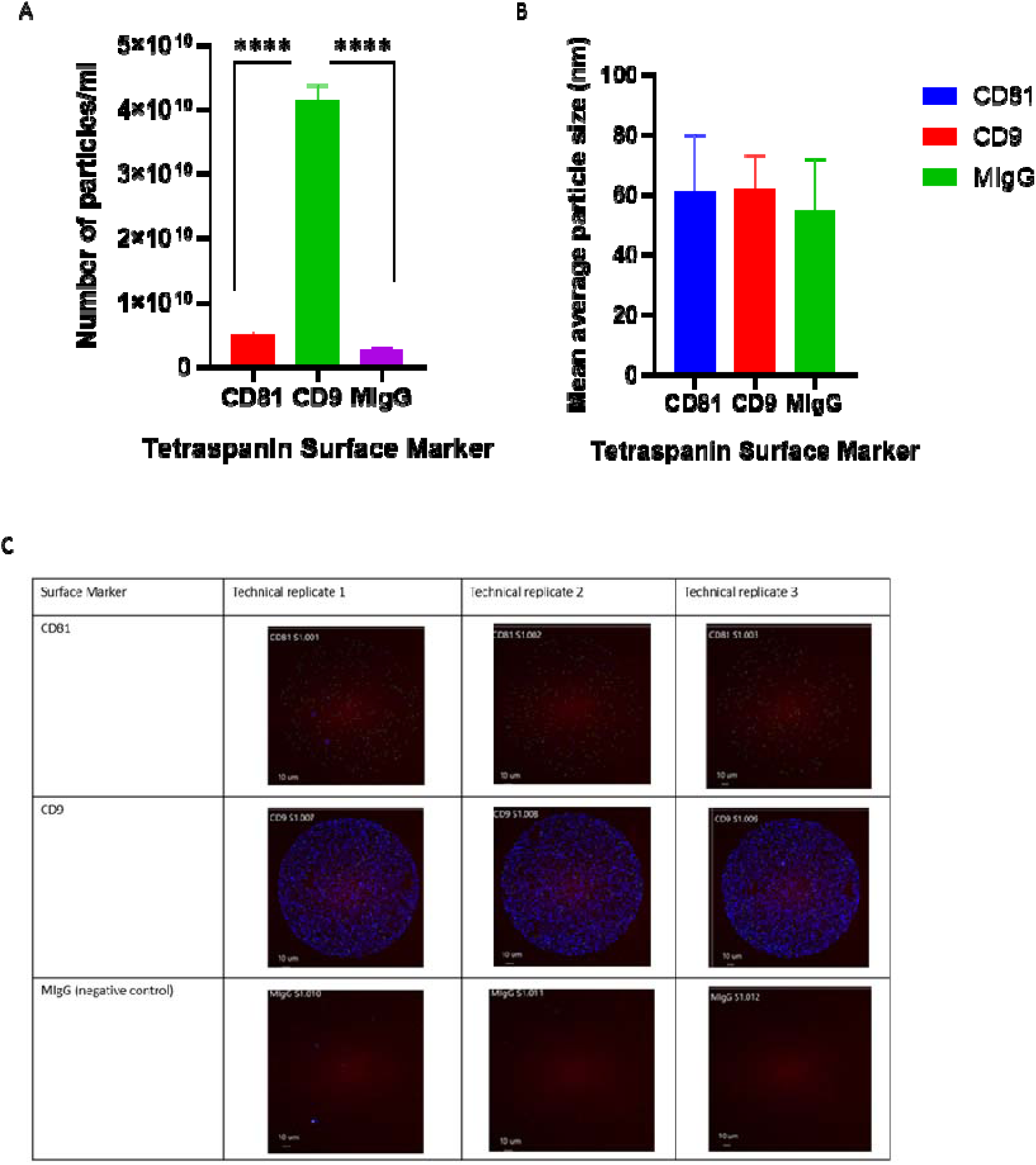
Exoview tetraspanin assay of PRP-EVs. (A) Particle concentration (particles/ml) per EV surface marker for PRP-EVs, with MIgG acting as a negative control, B) Mean average EV size per tetraspanin surface marker for PRP-EVs. Error bars are +/-standard deviation. (C) Visualization of PRP-EVs using Exoview Tetraspanin Assay. A fluorescent image of representative spots (technical replicates) is shown with colour denoting surface tetraspanin positive identification (blue-CD9 and green-CD81) Significance denoted by * (p<0.05), ** (p<0.01), *** (p< 0.001), **** (p<0.0001) following Shaprio Wilks normality testing and one-way ANOVA with multiple comparisons, followed by a post hoc Tukey test undertaken in GraphPad Prism.

### 3.2 PRP and PRP-EVs Proteomic Profiles

#### Distinct protein profiles were identified in PRP-EVs compared to matched plasma EVs and between PRP and plasma

We identified 575 proteins in PRP (22 unique to PRP), 558 in plasma (five unique to plasma), 209 in PRP-EVs (10 unique to PRP-EVs), 195 in plasma EVs (one unique protein found) (Figure 3A). There were 157 unique proteins (q <0.05) in PRP-EVs compared to plasma EVs, and 45 unique proteins in PRP compared with plasma. A Heatmap on quantified proteins following the removal of those with >50% missing values (Figure 3B), illustrated distinct differences in the proteome of based on EV source. This analysis was repeated comparing plasma to PRP (Figure 3C).

**Figure 3.**
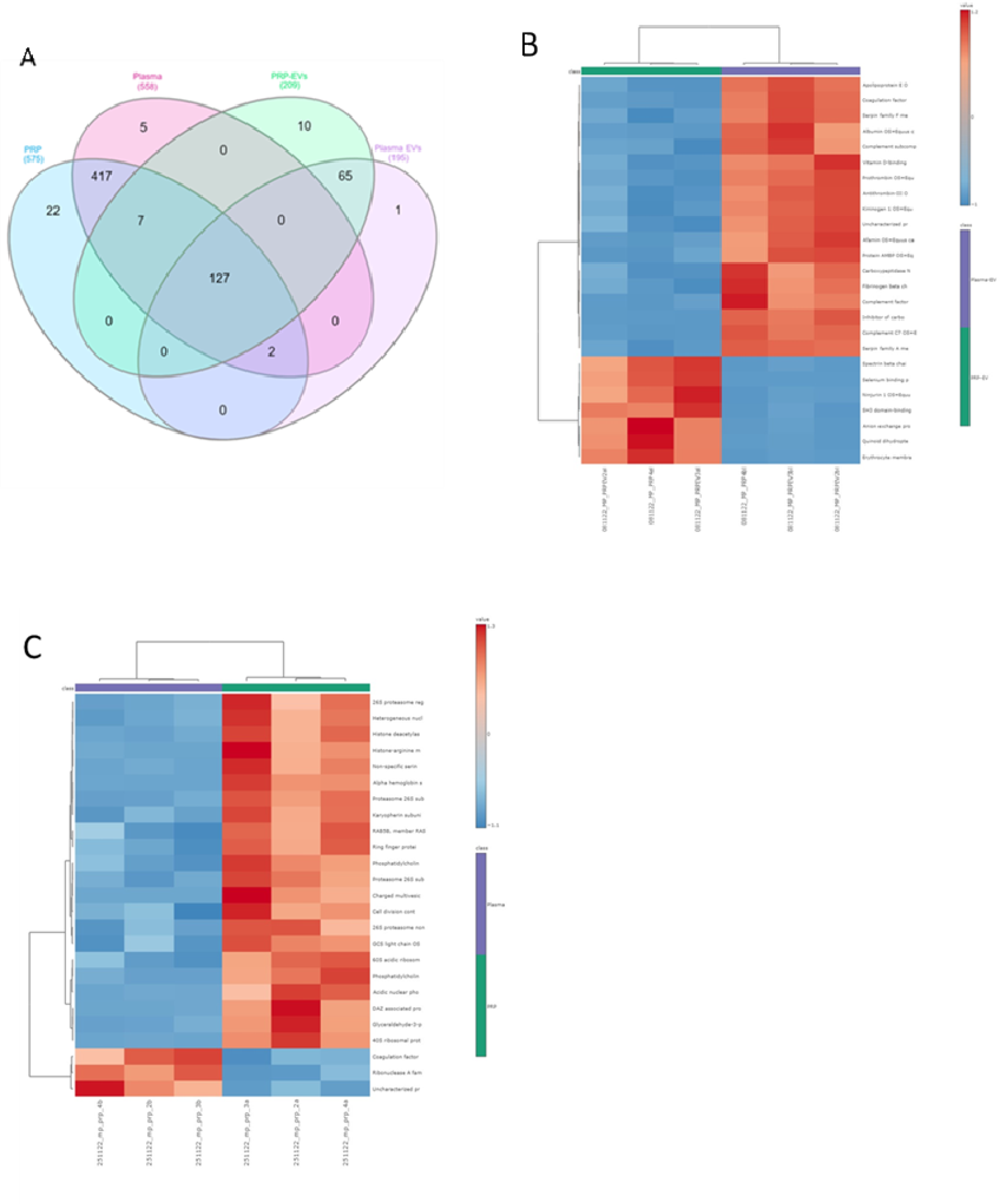
Proteins identified and quantified between PRP and PRP-EVs with matched plasma samples. (A) Venn diagram, (B) Heatmap analysis (using Euclidean distance and ward clustering) of the proteome of plasma-EVs and PRP-EVs (n=3) following Pareto scaling. (C) Heatmap analysis of the proteome of plasma and PRP (n=3). Data was analysed in Metaboanalyst v.5.0.

### 3.3 Functional Enrichment Analysis of PRP and PRP-EV Content

#### PRP was enriched with proteins associated with the regulation of cellular waste disposal whilst PRP-EVs showed a downregulation in lipid metabolism

Functional enrichment analysis was performed on differentially abundant (DA) proteins with and false discovery rate (FDR) p-value<0.05, and log fold change. The top five most significant canonical pathways for PRP versus plasma were: FAT10 signalling pathway (p=6.15×10^-4^), BAG2 signalling pathway (p=1.8×10^-3^), microautopahgy signalling pathway (p=4.37×10^-5^), inhibition of ARE mediated mRNA degradation pathway (p= 2.11×10^-3^) and protein ubiquitination pathway (p=1.38×10^-4^). Similarly, for PRP-EVs compared to plasma EVs, results were: acute phase response signalling (p=3.88×10^-5^), LXR-RXR activation (inhibited, p=1.02×10^-5^), FXR/RXR activation (p=1.59×10^-5^), DHCR24 signalling pathway (inhibited, p=9.29×10^-5^) and complement system (activated, p=1.39×10^-2^).

### 3.4 Gene Expression Analysis of In Vitro Tendon Inflammatory Model

#### Matrix metalloproteinases in tenocytes were elevated in the tendon inflammatory model, and there was a reduction in tendon marker expression

IGFBP6 was altered with respect to both treatment and cytokine stimulation (p=0.02). PRP and PRP-EV treatment did not restore IGFBP6 expression following pro-inflammatory stimulation. However, there was a significant increase in IGFBP6 when comparing no treatment and PRP treatment to EV treatment (p<0.01) (Figure 4A). COL1A2 expression changed with respect to both treatment (p<0.0001) and cytokine stimulation (p=0.0001). Increased expression in the control group following PRP-EV treatment, compared to no treatment and PRP treatment (q<0.0001) (Figure 4B) was observed. No increase in expression was observed for the cytokine stimulated groups following treatment. MMP13 expression was altered relative to both treatment and cytokine stimulation (p<0.0001). Within the cytokine stimulation group differences were observed between the no treatment and PRP-EV treatment (p<0.0001), and PRP treatment compared to PRP-EV treatment (p<0.0001); PRP treatment decreased expression compared to no treatment, and PRP-EV treatment resulted in MMP13 expression (Figure 4C). MMP3 expression increased following cytokine stimulation but did not reach significance. Following PRP treatment, MMP3 expression decreased, however following PRP-EV treatment MMP3 expression increased compared to control, though this did not reach statistical significance (Figure 4D). PRP treatment showed a trend towards decreasing MMP expression.

**Figure 4.**
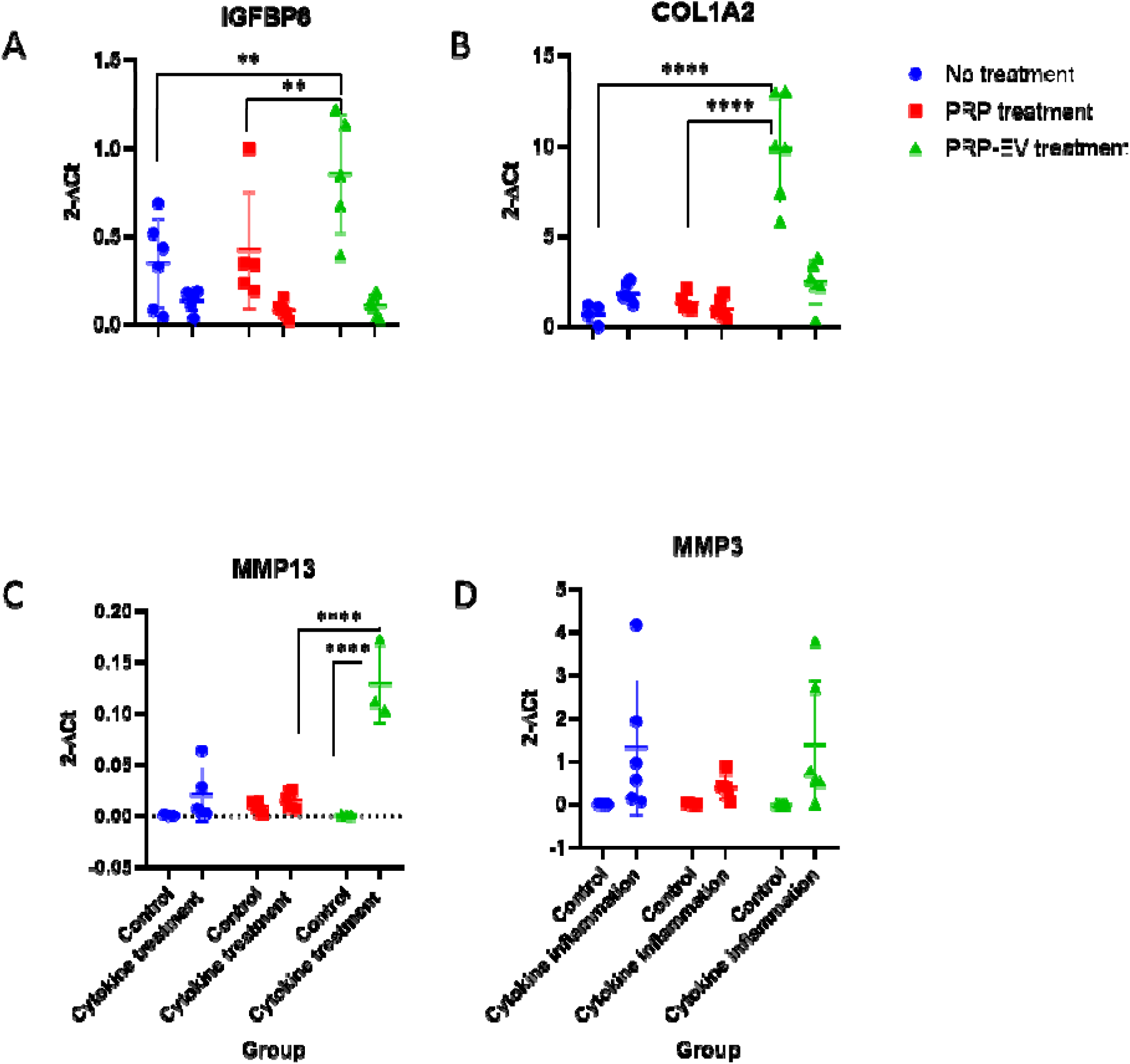
Gene expression analysis of control and cytokine stimulated equine tenocytes following PRP and PRP-EV treatments. A) IGFBP6, B) COL1A2, C) MMP13, D) MMP3 using RT-qPCR and relative quantification methods (2-Delta CT). Significance is denoted by an adjusted p value *(q<0.05), ** (q<0.01), *** (q< 0.001), **** (q<0.0001), following two-way ANOVA analysis and multiple comparison testing, using GraphPad Prism software v10.0 (California, USA).

### 3.5 Multiple Factor PCA

#### PRP and PRP-EVs altered the cellular proteome of cytokine stimulated equine tenocytes

A total of 1,933 proteins were identified across all samples and 1,567 proteins quantified. Following the removal of contaminating blood-derived proteins and proteins with 25% or more missing values a total of 832 proteins were used for downstream analysis. Multiple factor PCA of the proteome of all experimental groups was performed considering cellular phenotype (control or following cytokine stimulation) and the effect of treatment (no treatment, PRP treatment and PRP-EV treatment). There was a change following PRP or PRP-EV treatment (Figure 5) with a shift towards that of control tenocytes when cells were treated with PRP or PRP-EVs following cytokine treatment.

**Figure 5.**
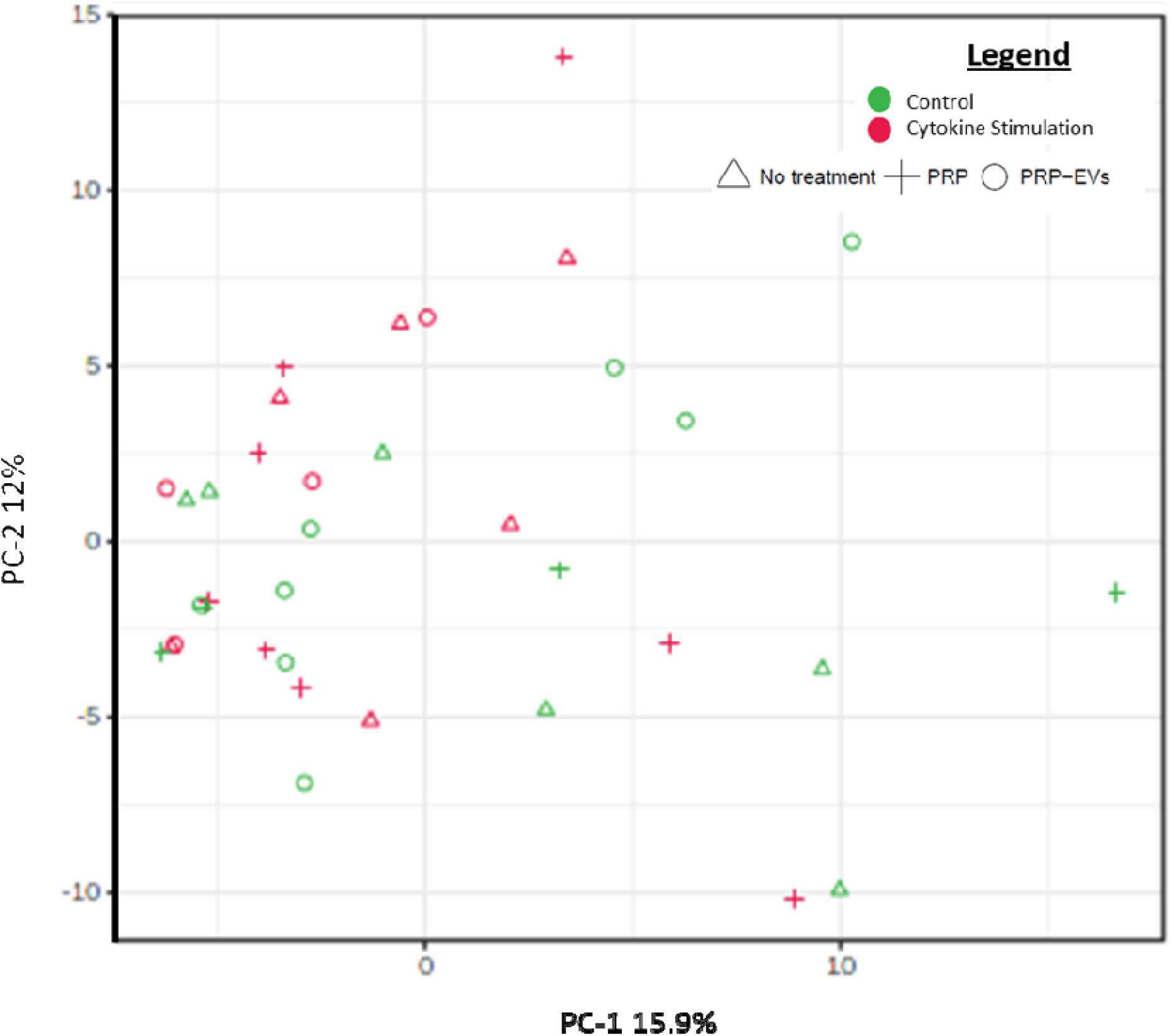
Unsupervised multivariate analysis using principal component analysis. Red denotes cytokine treatment and green denotes control, a triangle symbolises n treatment, a cross represents samples treated with PRP and a circle represents PRP-EV treatment.

*3.6 Cellular Proteome Analysis*

#### 18 proteins changed following cytokine stimulation and treatment of either platelet rich plasma or platelet rich plasma derived extracellular vesicles

A two-way ANOVA was performed on all cellular proteomics data to account for both categorical variables within the study: cytokine stimulation (control or cytokine stimulation) and treatment (no treatment, PRP or PRP-EVs) and their effect on protein expression (Figure 6). 18 proteins were identified as significant with respect to both categoric variables at an unadjusted p<0.05. These proteins included the following with p values corresponding to the effect of cytokine stimulation and PRP or PRP-EV treatment, respectively: adenylate kinase 2 mitochondrial protein (p=0.035 and p=0.038) (Supplementary Figure 1A), AHNAK nucleoprotein (p=7.19E-5 and p=0.01) (Figure 6A), Beta-galactosidase (p=0.028 and p=0.046), centrosomal protein of 162kDa (p=0.013 and p=0.019), col1a1 chain (p=4.63E-5 and p=0.012) (Figure 6F), D-3-phosphoglycerate dehydrogenase (p=0.047 and p=0.034), F-actin-capping protein subunit alpha (p=0.042 and p=0.02), heme oxygenase (p= 0.002 and p=0.004) (Figure 6E), heterogeneous nuclear ribonucleoprotein U like 1 (p=0.011 and p=0.035), LRP chaperone MESD (p=0.038 and p=0.043), polyadenylate-binding protein (p=0.022 and p=0.009), protein disulphide isomerase (p=0.022 and p= 0.012), SAP domain containing ribonucleoprotein (p= 0.021 and p=0.003) (Figure 6B), sequestosome 1 (p=0.001 and p=0.036) (Figure 6C), small monomeric GTPase (p=0.017 and p=0.036), sulphide quinone oxidoreductase (p=0.022 and p=0.022), THADA armadillo repeat (p=0.02 and p= 0.012) and translocon associated protein subunit alpha (p=0.003 and p=0.01) (Figure 6D). P values were subsequently used in pathway analysis as it has been reported that multiple testing correction methods can fail to identify statistically significant values due to stringent thresholds 51. Figure 6 depicts a selection of significant proteins identified following two-way ANOVA and post hoc Tukey test. Figure 6A demonstrates reduced expression of AHNAK nucleoprotein between control and cytokine treatment following PRP treatment (p<0.01). In addition, there was a significant difference in protein abundance between PRP and PRP-EV treatment (p<0.05). Figure 6B demonstrates a significant increase in SAP domain containing ribonucleoprotein between control and the cytokine following PRP treatment (p<0.01), and a difference in abundance between no treatment and PRP treatment (p<0.05). Figure 6C highlights a significant increase in sequestosome 1 between control and the cytokine following PRP and PRP-EV treatment (p<0.05). In addition, a difference in protein abundance between no treatment and PRP-EV treatment groups (p<0.05) was evident. Figure 6D shows a significant decrease in translocon associated protein subunit alpha expression between control and cytokine following PRP treatment (p<0.05) and a difference between no treatment and PRP treatment (p<0.05). Figure 6E highlights a significant increase in heme oxygenase following cytokine treatment when compared to control (p<0.01), and a difference between no treatment and PRP and PRP-EV treatment groups (p<0.05). Finally, Figure 6F shows a reduction in col1a1 abundance between control and cytokine (p<0.05), and between control and cytokine treated with PRP (p<0.01) in both the control and PRP treated cytokine groups. It was also different between the control and that of PRP-EV treatment (p<0.05).

**Figure 6.**
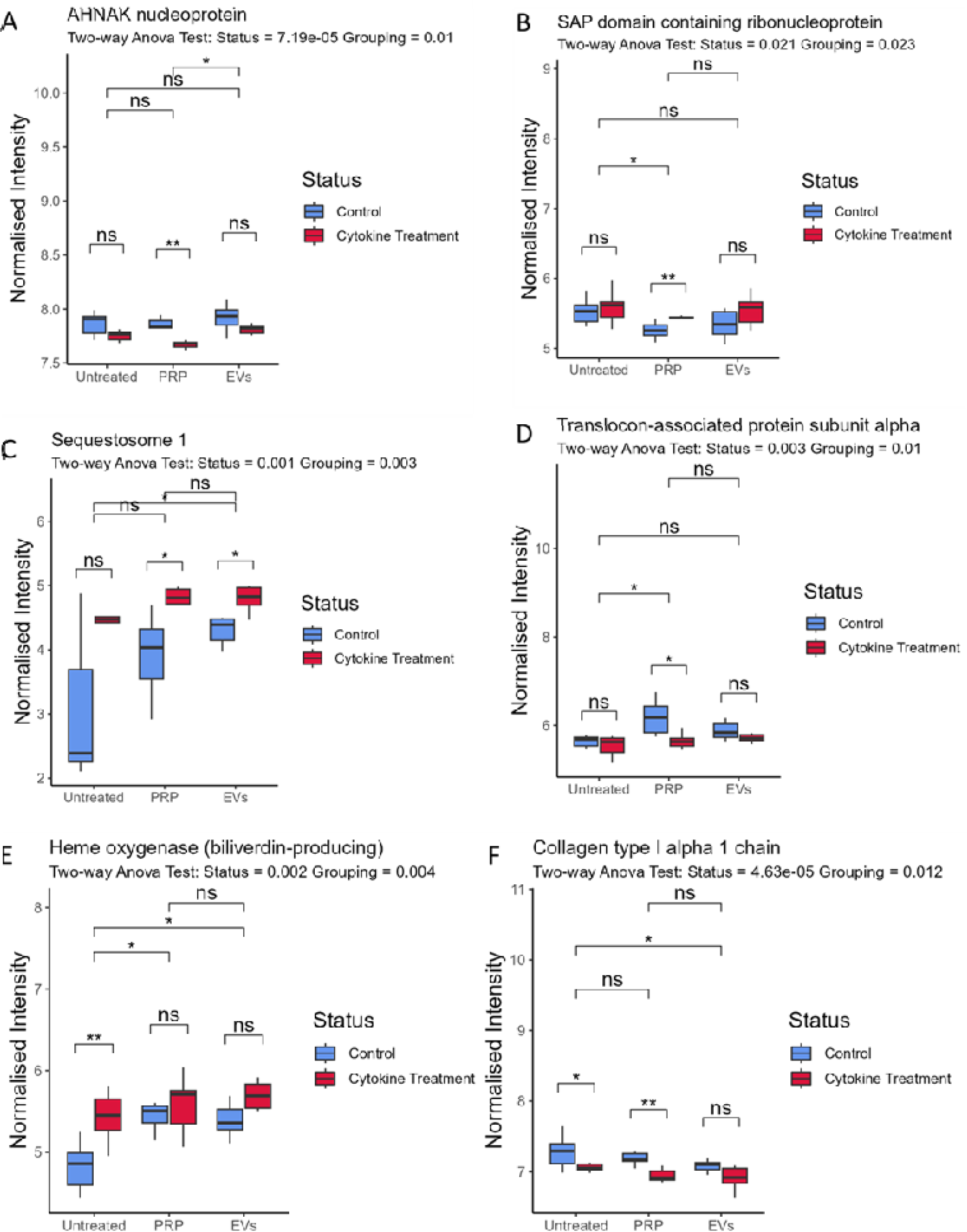
Differential abundant proteins following cytokine stimulation of equine tenocytes. The effect of no treatment, PRP and PRP-EV treatment, as identified by a two-way ANOVA and post hoc tukey test; A) AHNAK nucleoprotein, (B) SAP domain containing ribonucleoprotein, (C) sequestosome 1, (D) translocon associated protein subunit alpha, (E) heme oxygenase and (F) col1a1. Significance is denoted by *(p<0.05), ** (p<0.01), *** (p< 0.001), **** (p<0.0001). Statistical analysis and graphs made using R studio software.

## 4. Discussion

Our pilot study determined the effect of PRP and PRP-EVs on tendon inflammation using an *in vitro* model. To our knowledge this is the first study of its kind to explore the role of PRP-EVs in PRP therapeutic mediation *in vitro*. The rationale for determining the cargo of PRP-EVs was that others had demonstrated that platelets secrete a significant number of EVs^27^. We hypothesised that EVs may be involved in the therapeutic action of PRP.

PRP-EVs are a heterogenous population of EVs from multiple sources, including leukocytes (T-cells and B cells) ^52^ and an enrichment in platelet derived EVs. Pre-clinical studies have evidenced the potential of PRP-EVs in future therapeutics, due to their size and stability ^28^. We found that PRP and PRP-EVs were enriched in proteins associated with cellular waste disposal and lipid metabolism, respectively when compared to plasma and plasma-EVs. Whilst not significant our results suggest there may be a reduction in the expression of matrix gene expression following PRP treatment. Proteomics revealed that PRP and PRP-EV treatment had a small effect on control equine tenocytes and those treated with cytokines (18 significant proteins across cytokine stimulation and treatment; no treatment, PRP, PRP-EVs). However, these proteins appeared to be involved in pathways associated with collagen metabolism and NFkB signalling. As proteomic differences between PRP and PRP-EV treatment were not significant globally this may be suggestive of PRP-EVs role in mediating PRP therapeutic action.

We interrogated the PRP and PRP-EV proteomes to determine significant pathways. It was found that enriched pathways in PRP included microautophagy signalling pathway and the protein ubiquitination pathways. Microautopahgy is a non-selective lysosomal degradative process, which engulfs cytoplasmic cargo at membrane boundaries via autophagic tubes ^53,54^. In this study proteins including proteasome 26s subunit ATPase 1, 2, 3, 4,5 and 6 were mapped to this pathway. Previous studies have implicated autophagy more broadly in the development and repair of tendon following a traumatic injury55. Autophagy prevented the loss of stemness in tendon stem cells as a result of increased oxidative stress often evident at the site of tendon injury, by supressing the accumulation of reactive oxygen species (ROS) ^56^. Thus, PRP may have the capacity to enhance this pathway, acting in a protective anti-oxidative capacity, protecting resident cells from damage. Protein ubiquitination is involved in the degradation of over 80% of proteins in cells and is regarded as a reversible post-translational modification. In a rotator cuff injury study, the ubiquitin-proteasome pathway was a potent regulator of muscular atrophy and thus protein degradation can be attributed to such occurrence as a result of tendon injury ^57^. PRP was enriched in proteins that can activate autophagic and ubiquitination processes, potentially promoting collagen metabolism, skeletal tissue homeostasis and reducing oxidative stress. This could have significant implications when considering the repair of tendon injury sites. Following functional enrichment analysis of the proteome of PRP-EVs we identified acute phase response signalling (inhibited), LXR-RXR (inhibited) and FXR-RXR signalling (not stated) as canonical pathways associated with PRP-EV cargo. Acute phase response signalling was associated with the enhancement of late-stage healing in tendon injury, induced by IL-6, despite the role of IL-6 in inflammation and cellular response to tissue injury 12. It was directly attributed to inflammation due to cytokine regulation and involved multiple plasma proteins including C-reactive protein, serum amyloid A, transthyretin, fibronectin^58^. This suggests that PRP-EVs could-potentially act in an anti-inflammatory manner by inhibiting acute phase response signalling. LXR-RXR signalling was predicted to be inhibited in this study. LXRs are transcriptional regulators of lipid homeostasis, with anti-inflammatory properties ^59,60^. RXR is a nuclear hormone receptor of the retinoid family, that functions in conjunction with FXR and LXR. FXR-RXR signalling is involved in the activation of cholesterol homeostasis, and suppression of inflammatory processes ^60^. In this study both LXR and FXR – RXR signalling were attributed to PRP-EV cargo. This suggests that PRP-EVs cargo could inhibit anti-inflammatory mechanisms. PRP can elicit an inflammatory response in human tendon fibroblasts *in vitro*, by stimulating TNF-α and NFkB pathways, resulting in the formation of ROS and the activation of oxidative stress pathways ^61^. Therefore, PRP may act by causing an acute inflammatory event that stimulates endogenous tissue regeneration responses ^61^, but further research is required to elucidate definitive pro or anti-inflammatory activity, especially for PRP-EVs.

We found the tendon marker IGFBP6 ^62^ was significantly downregulated following cytokine treatment which is characteristic of an impaired tendon phenotype ^63^.

A global proteome shift was observed using multiple factor PCA in response to cytokine stimulation. Further analysis elucidated potential proteomic drivers of this variation. 18 proteins were significant with respect to both cytokine stimulation and treatment. One of these proteins, sequestosome 1 acts as a cargo receptor and is involved in the degradation of ubiquitinated proteins from both autophagic ad proteasomal pathways ^64^. It is a regulator of NFkB signalling ^65^ which is important in inflammation, cellular stress response and cellular survival ^66^. NFkB is a transcription factor, stimulated by pro-inflammatory cytokines, chemokines, stress-related factors and ECM degradation products. Upon stimulation NFkB molecules can trigger the expression of genes associated with tissue destruction ^67^. Previous studies have implicated NFkB pathway signalling in tendinopathy. In murine tendon fibroblasts with cre-mediated overexpression of IKb, there was increased degeneration of mouse rotator cuff tendons which corresponded to increased levels of proinflammatory cytokines and innate immune cells within the joint ^68^. Targeting of the NFkB pathway may serve as a prospective therapeutic approach in tendon disorders, and subsequent expression of sequestosome 1 would serve as a marker of this. Our study demonstrated that cytokine stimulation increased sequestosome 1 expression and that addition of PRP did not significantly change it. However, addition of PRP-EVs did. Thus, the EV component of PRP may elicit a localised inflammatory response, whereas other biological factors in PRP may prevent as great an effect occurring. Col1a1 was differentially abundant across experimental groups with respect to global proteome analysis. Type 1 collagen is the predominant tendon collagen ^69^ but expression was decreased during the healing process following tendon injury ^70,71^. Over time some of the granulated tissue is replaced with type 1 collagen. Subsequently type 3 collagen is associated with scar tissue following injury. In addition, an increase in the type 3/ type 1 collagen ratio was observed with age ^72^ and pathology ^73,74^. Upon cytokine stimulation in our study tenocyte abundance of col1a1 decreased. PRP treatment did not alter col1a1 abundance compared to the non-treatment group, with col1a1 having a significantly decreased abundance in the cytokine stimulation group compared to control, irrespective of treatment. This suggests PRP may not act on collagen metabolism. However, PRP-EV treatment of cytokine stimulated tenocytes did cause differential abundance of col1a1 when compared with untreated groups, and no significant difference was observed between control and cytokine stimulation following PRP-EV treatment. Potentially PRP-EVs may alter collagen production to baseline and components of PRP (platelets and plasma) may contain biological agents that reduce this effect. Hence to understand this relationship further mechanistic studies are required to ascertain if PRP contains inhibitory biological agents with implications for therapeutic efficacy.

There were limitations associated with this study including the low sample size. As a result, p values were used when analysing MS proteomic data from the *in vitro* experiment. This is however common practice in proteomic studies as multiple testing correction methods can fail to identify statistically significant values due to too stringent threshold in the case of underpowered studies ^51^. Moreover, the use of monolayer culture can alter the physiological outcomes observed in tendon injury-based studies, as they do not account for the three-dimensional environment tenocytes are situated within and their role in responding to external stimuli and biomechanical forces ^75,76^. Tenocytes in monolayer culture can be unstable phenotypically and dedifferentiate ^77^. Future studies should explore the use of bioengineering approaches such as microfluidic technologies ^78^. In addition, there were variables that could not be accounted for within this study that may affect reported outcomes, such as donor variation in EV uptake and response to PRP. Additional studies could use bioluminescent or fluorescent labelling approaches for imaging to visualise cellular uptake of EVs ^79^. Finally, allogenic PRP and subsequent PRP-EV treatment was used rather than the traditional autologous PRP product utilised in clinical practice. This may have elicited a different cellular immune response compared to tendon cell culture in which no white blood cells are present.

## 5. Conclusion

This *in vitro* pilot study identified potential molecular mechanisms mediating the therapeutic effect observed in vivo. These require further investigation and validation, particularly around the role of collagen metabolism and NF-kB signalling in association with inflammation. Evidence was provided that PRP-EVs elicit an effect on stimulated tenocytes and thus may be involved in the mediation of the therapeutic product, and at a minimum should be examined further when standardising PRP products. We highlight initial evidence that an EV based product could be available ‘off the shelf’ to treat tendon injury.

## 6. Ethics

Ethical approval was obtained from the University of Liverpool Veterinary Research Ethics Committee. Tenocyte samples from both our equine musculoskeletal biobank (VREC561) and clinical donors from the Philip Leverhulme equine hospital were used within this study (VREC561). Venous whole blood was collected from an abattoir (VREC561).

## References

1. Thorpe CT, Clegg PD, Birch HL. A review of tendon injury: Why is the equine superficial digital flexor tendon most at risk? Equine Vet J. 2010;42(2):174–180. doi:10.2746/042516409X480395

2. Murray RC, Dyson SJ, Tranquille C, Adams V. Association of type of sport and performance level with anatomical site of orthopaedic injury diagnosis. Equine Vet J. 2006;38(SUPPL.36):411–416. doi:10.1111/j.2042-3306.2006.tb05578.x

3. O’Brien C, Marr N, Thorpe C. Microdamage in the equine superficial digital flexor tendon. Equine Vet J. 2021;53(3):417–430. doi:10.1111/evj.13331

4. Patterson-Kane JC, Becker DL, Rich T. The Pathogenesis of Tendon Microdamage in Athletes: The Horse as a Natural Model for Basic Cellular Research. J Comp Pathol. 2012;147(2-3):227–247. doi:10.1016/j.jcpa.2012.05.010

5. Tamura N, Kodaira K, Yoshihara E, … NMTV, 2018 undefined. A retrospective cohort study investigating risk factors for the failure of Thoroughbred racehorses to return to racing after superficial digital flexor tendon injury. Elsevier.

6. Perkins NR, Reid SWJ, Morris RS. Risk factors for injury to the superficial digital flexor tendon and suspensory apparatus in Thoroughbred racehorses in New Zealand. N Z Vet J. 2005;53(3):184–192. doi:10.1080/00480169.2005.36503

7. Lam KH, Parkin TDH, Riggs CM, Morgan KL. Descriptive analysis of retirement of Thoroughbred racehorses due to tendon injuries at the Hong Kong Jockey Club (1992-2004). Equine Vet J. 2007;39(2):143–148. doi:10.2746/042516407X159132

8. Lam KKH, Parkin TDH, Riggs CM, Morgan KL. Evaluation of detailed training data to identify risk factors for retirement because of tendon injuries in Thoroughbred racehorses. Am J Vet Res. 2007;68(11):1188–1197. doi:10.2460/ajvr.68.11.1188

9. Marr CM, Mcmillant I, Boyd JS, Wright NG, Murray M. U ltrasonograp h ic and histopat holog ical findings in equine superficial digital flexor tendon injury. 25(1993):23-29.

10. Johnson SA, Valdés-Martínez A, Turk PJ, et al. Longitudinal tendon healing assessed with multi-modality advanced imaging and tissue analysis. Equine Vet J. 2021;(January):1–16. doi:10.1111/evj.13478

11. Thorpe CT, Clegg PD, Birch HL. A review of tendon injury: Why is the equine superficial digital flexor tendon most at risk? Equine Vet J. 2010;42(2):174–180. doi:10.2746/042516409X480395

12. Chisari E, Rehak L, Khan WS, Maffulli N. Tendon healing is adversely affected by low-grade inflammation. J Orthop Surg Res. 2021;16(1):1–9. doi:10.1186/s13018-021-02811-w

13. Del Buono A, Battery L, Denaro V, Maccauro G, Maffulli N. Tendinopathy and inflammation: some truths. Int J Immunopathol Pharmacol. 2011;24(1 Suppl 2):45–50. doi:10.1177/03946320110241s209

14. Beerts C, Suls M, Broeckx SY, et al. Tenogenically induced allogeneic peripheral blood mesenchymal stem cells in allogeneic platelet-rich plasma: 2-year follow-up after tendon or ligament treatment in horses. Front Vet Sci. 2017;4(SEP). doi:10.3389/fvets.2017.00158

15. Brossi PM, Moreira JJ, Machado TSL, Baccarin RYA. Platelet-rich plasma in orthopedic therapy: A comparative systematic review of clinical and experimental data in equine and human musculoskeletal lesions. BMC Vet Res. 2015;11(1). doi:10.1186/S12917-015-0403-Z

16. Wijekoon HMS, de Silva DDN. Current Evidence on Using Platelet Rich Plasma as a Therapeutic Modality for Veterinary Orthopedic Conditions. World’s Veterinary Journal. 2021;11(1):73–78. doi:10.54203/scil.2021.wvj10

17. Bosch G, Moleman M, Barneveld A, van Weeren PR, van Schie HTM. The effect of platelet-rich plasma on the neovascularization of surgically created equine superficial digital flexor tendon lesions. Scand J Med Sci Sports. 2011;21(4):554–561. doi:10.1111/j.1600-0838.2009.01070.x

18. Parrish WR, Roides B, Hwang J, Mafilios M, Story B, Bhattacharyya S. Normal platelet function in platelet concentrates requires non-platelet cells: A comparative in vitro evaluation of leucocyte-rich (type 1a) and leucocyte-poor (type 3b) platelet concentrates. BMJ Open Sport Exerc Med. 2016;2(1):1–8. doi:10.1136/bmjsem-2015-000071

19. Fortier L. Clinical Use of Stem Cells, Marrow Components, and Other Growth Factors. Second Edi. Elsevier Inc.; 2010. doi:10.1016/B978-1-4160-6069-7.00073-0

20. Geburek F, Gaus M, van Schie HTM, Rohn K, Stadler PM. Effect of intralesional platelet-rich plasma (PRP) treatment on clinical and ultrasonographic parameters in equine naturally occurring superficial digital flexor tendinopathies - a randomized prospective controlled clinical trial. BMC Vet Res. 2016;12(1). doi:10.1186/s12917-016-0826-1

21. Bosch G, Van Schie HTM, De Groot MW, et al. Effects of platelet-rich plasma on the quality of repair of mechanically induced core lesions in equine superficial digital flexor tendons: A placebo-controlled experimental study. Journal of Orthopaedic Research. 2010;28(2):211–217. doi:10.1002/jor.20980

22. Montano C, Auletta L, Greco A, et al. The Use of Platelet-Rich Plasma for Treatment of Tenodesmic Lesions in Horses: A Systematic Review and Meta-Analysis of Clinical and Experimental Data. mdpi.com. Published online 2021. doi:10.3390/ani11030793

23. Wu J, Piao Y, Liu Q, Yang X. Platelet-rich plasma-derived extracellular vesicles: A superior alternative in regenerative medicine? Cell Prolif. 2021;54(12). doi:10.1111/cpr.13123

24. Zakirova EY, Aimaletdinov AM, Malanyeva AG, Rutland CS, Rizvanov AA. Extracellular Vesicles: New Perspectives of Regenerative and Reproductive Veterinary Medicine. Front Vet Sci. 2020;7(November):1–5. doi:10.3389/fvets.2020.594044

25. Graça AL, Domingues RMA, Calejo I, Gómez-Florit M, Gomes ME. Therapeutic Effects of Platelet-Derived Extracellular Vesicles in a Bioengineered Tendon Disease Model. Int J Mol Sci. 2022;23(6). doi:10.3390/ijms23062948

26. Puhm F, Boilard E, MacHlus KR. Platelet extracellular vesicles; beyond the blood. Arterioscler Thromb Vasc Biol. 2021;41(1):87–96. doi:10.1161/ATVBAHA.120.314644

27. Tao SC, Guo SC, Zhang CQ. Platelet-derived Extracellular Vesicles: An Emerging Therapeutic Approach. Int J Biol Sci. 2017;13(7):828–834. doi:10.7150/ijbs.19776

28. Wu J, Piao Y, Liu Q, Yang X. Platelet-rich plasma-derived extracellular vesicles: A superior alternative in regenerative medicine? Cell Prolif. 2021;54(12):1–13. doi:10.1111/cpr.13123

29. Clarke, Lima C, Anderson J, et al. Optical photothermal infrared spectroscopy can differentiate equine osteoarthritic plasma extracellular vesicles from healthy controls. Analytical Methods. Published online 2022:2022.03.11.483922. doi:10.1101/2022.03.11.483922

30. Clarke, Johnson, Gutierrez C, et al. Temporal extracellular vesicle protein changes following intraarticular treatment with integrin α10β1-selected mesenchymal stem cells in equine osteoarthritis. Front Vet Sci. 2022;9. doi:10.3389/fvets.2022.1057667

31. Turlo AJ, Ashraf Kharaz Y, Clegg PD, Anderson J, Peffers MJ. Donor age affects proteome composition of tenocyte-derived engineered tendon. BMC Biotechnol. 2018;18(1):1–11. doi:10.1186/s12896-018-0414-5

32. Peffers MJ, Liu X, Clegg PD. Transcriptomic signatures in cartilage ageing. Arthritis Res Ther. 2013;15(4). doi:10.1186/ar4278

33. Chomczynski P, Sacchi N. Single-step method of RNA isolation by acid guanidinium thiocyanate-phenol-chloroform extraction. Anal Biochem. 1987;162(1):156–159. doi:10.1016/0003-2697(87)90021-2

34. Peffers MJ, Liu X, Clegg PD. Transcriptomic signatures in cartilage ageing. Arthritis Res Ther. 2013;15(4). doi:10.1186/ar4278

35. Livak KJ, Schmittgen TD. Analysis of relative gene expression data using real-time quantitative PCR and the 2-ΔΔCT method. Methods. 2001;25(4):402–408. doi:10.1006/meth.2001.1262

36. Ragni E, Orfei CP, Bowles AC, Girolamo L De, Correa D. Reliable reference genes for gene expression assessment in tendon-derived cells under inflammatory and pro-fibrotic/healing stimuli. Cells. 2019;8(10):1–12. doi:10.3390/cells8101188

37. Peffers MJ, Collins J, Loughlin J, Proctor C, Clegg PD. A proteomic analysis of chondrogenic, osteogenic and tenogenic constructs from ageing mesenchymal stem cells. Stem Cell Research and Therapy. 2016;7(1):1–16. doi:10.1186/s13287-016-0384-2

38. Peffers MJ, Thorpe CT, Collins JA, et al. Proteomic analysis reveals age-related changes in tendon matrix composition, with age- and injury-specific matrix fragmentation. Journal of Biological Chemistry. 2014;289(37):25867–25878. doi:10.1074/jbc.M114.566554

39. Peffers MJ, Thorpe CT, Collins JA, et al. Proteomic Analysis Reveals Age-related Changes in Tendon Matrix Composition, with Age- and Injury-specific Matrix Fragmentation. Journal of Biological Chemistry. 2014;289(37):25867–25878. doi:10.1074/jbc.M114.566554

40. Peffers MJ, Collins J, Loughlin J, Proctor C, Clegg PD. A proteomic analysis of chondrogenic, osteogenic and tenogenic constructs from ageing mesenchymal stem cells. Stem Cell Res Ther. 2016;7(1):133. doi:10.1186/s13287-016-0384-2

41. Anderson, Phelan, Foddy, Clegg, Peffers. Ex Vivo Equine Cartilage Explant Osteoarthritis Model: A Metabolomics and Proteomics Study. ACS Publications. 2020;19(9):3652–3667. doi:10.1021/acs.jproteome.0c00143

42. Anderson, Phelan, Rubio-Martinez, et al. Optimization of Synovial Fluid Collection and Processing for NMR Metabolomics and LC-MS/MS Proteomics. J Proteome Res. 2020;19(7):2585–2597. doi:10.1021/acs.jproteome.0c00035

43. Anderson JR, Smagul A, Simpson D, Clegg PD, Rubio-Martinez LM, Peffers MJ. The synovial fluid proteome differentiates between septic and nonseptic articular pathologies. J Proteomics. 2019;202(January). doi:10.1016/j.jprot.2019.04.020

44. Peffers M, Riggs C, McDermott B, Clegg P. Comprehensive Protein Profiling of Synovial Fluid in Osteoarthritis. Equine Vet J. 2014;46:19–19. doi:10.1111/evj.12323_42

45. Anderson JR, Phelan MM, Rubio-Martinez LM, et al. Optimization of Synovial Fluid Collection and Processing for NMR Metabolomics and LC-MS/MS Proteomics. J Proteome Res. 2020;19(7):2585–2597. doi:10.1021/acs.jproteome.0c00035

46. Anderson, Phelan, Foddy, Clegg, Peffers. Ex Vivo Equine Cartilage Explant Osteoarthritis Model: A Metabolomics and Proteomics Study. J Proteome Res. 2020;19(9):3652–3667. doi:10.1021/acs.jproteome.0c00143

47. Ashraf Kharaz Y, Zamboulis D, Sanders K, Comerford E, Clegg P, Peffers M. Comparison between chaotropic and detergent-based sample preparation workflow in tendon for mass spectrometry analysis. Proteomics. 2017;17(13-14):13–14. doi:10.1002/pmic.201700018

48. Perez-Riverol Y, Csordas A, … JB. The PRIDE database and related tools and resources in 2019: improving support for quantification data. Nucleic Acid Res. Published online 2019.

49. Pang Z, Chong J, Zhou G, et al. MetaboAnalyst 5.0: narrowing the gap between raw spectra and functional insights. Nucleic Acids Res. 2021;49(W1):W388–W396. doi:10.1093/nar/gkab382

50. Clarke EJ, Johnson E, Caamaño Gutierrez E, et al. Temporal extracellular vesicle protein changes following intraarticular treatment with integrin α10β1-selected mesenchymal stem cells in equine osteoarthritis. Front Vet Sci. 2022;9. doi:10.3389/fvets.2022.1057667

51. Pascovici D, Handler DCL, Wu JX, Haynes PA. Multiple testing corrections in quantitative proteomics: A useful but blunt tool. Proteomics. 2016;16(18):2448–2453. doi:10.1002/pmic.201600044

52. Auber M, Svenningsen P. An estimate of extracellular vesicle secretion rates of human blood cells. Journal of Extracellular Biology. 2022;1(6). doi:10.1002/jex2.46

53. Li WW, Li J, Bao JK. Microautophagy: Lesser-known self-eating. Cellular and Molecular Life Sciences. 2012;69(7):1125-1136. doi:10.1007/s00018-011-0865-5

54. Kawamura N, Sun-Wada GH, Aoyama M, et al. Delivery of endosomes to lysosomes via microautophagy in the visceral endoderm of mouse embryos. Nat Commun. 2012;3(May). doi:10.1038/ncomms2069

55. Montagna C, Svensson RB, Bayer ML, et al. Autophagy guards tendon homeostasis. Cell Death Dis. 2022;13(4). doi:10.1038/s41419-022-04824-7

56. Chen H, Ge HA, Wu GB, Cheng B, Lu Y, Jiang C. Autophagy Prevents Oxidative Stress-Induced Loss of Self-Renewal Capacity and Stemness in Human Tendon Stem Cells by Reducing ROS Accumulation. Cellular Physiology and Biochemistry. 2016;39(6):2227–2238. doi:10.1159/000447916

57. Sunil K, Joshi B, Hubert T, Kim MD, Brian T, Xuhui L. Differential ubiquitin-proteasome and autophagy signaling following rotator cuff tears and suprascapular nerve injury. Journal of Orthopaedic Research. Published online 2014. doi:10.1002/jor.22482.Differential

58. Miroshnychenko O, Chalkley RJ, Leib RD, Everts PA, Dragoo JL. Proteomic analysis of platelet-rich and platelet-poor plasma. Regen Ther. 2020;15:226–235. doi:10.1016/j.reth.2020.09.004

59. Ito A, Hong C, Rong X, et al. LXRs link metabolism to inflammation through Abca1-dependent regulation of membrane composition and TLR signaling. Elife. 2015;4(JULY 2015):1–23. doi:10.7554/eLife.08009

60. Kong Z, Zhou C, Chen L, et al. Multi-omics analysis reveals up-regulation of APR signaling, LXR/RXR and FXR/RXR activation pathways in holstein dairy cows exposed to high-altitude hypoxia. Animals. 2019;9(7):1–15. doi:10.3390/ani9070406

61. Hudgens JL, Sugg KB, Grekin JA, Gumucio JP, Bedi A, Mendias CL. Platelet-Rich Plasma Activates Proinflammatory Signaling Pathways and Induces Oxidative Stress in Tendon Fibroblasts. American Journal of Sports Medicine. 2016;44(8):1931–1940. doi:10.1177/0363546516637176

62. Turlo AJ, Mueller-Breckenridge AJ, Zamboulis DE, Tew SR, Canty-Laird EG, Clegg PD. Insulin-like growth factor binding protein (Igfbp6) is a cross-species tendon marker. Eur Cell Mater. 2019;38:123–136. doi:10.22203/eCM.v038a10

63. Dahlgren LA, Mohammed HO, Nixon AJ. Expression of insulin-like growth factor binding proteins in healing tendon lesions. Journal of Orthopaedic Research. 2006;24(2):183–192. doi:10.1002/jor.20000

64. Philippe C. Linking Amyotrophic Lateral Sclerosis and Frontotemporal Dementia. Elsevier Inc.; 2020. doi:10.1016/B978-0-12-815854-8.00004-5

65. Ratti E, Berry JD. Amyotrophic Lateral Sclerosis 1 and Many Diseases. Elsevier Inc.; 2016. doi:10.1016/B978-0-12-800105-9.00042-1

66. Rea SL, Walsh JP, Layfield R, Ratajczak T, Xu Jiake J. New insights into the role of sequestosome 1/p62 mutant proteins in the pathogenesis of paget’s disease of bone. Endocr Rev. 2013;34(4):501–524. doi:10.1210/er.2012-1034

67. Rigoglou S, Papavassiliou AG. The NF-κB signalling pathway in osteoarthritis. International Journal of Biochemistry and Cell Biology. 2013;45(11):2580–2584. doi:10.1016/j.biocel.2013.08.018

68. Abraham A, Shah S, Golman M, et al. Targeting the NF-κB signaling pathway in chronic tendon disease. Sci Transl Med. 2019;11(481). doi:10.1126/scitranslmed.aav4319.Targeting

69. Salvatore L, Gallo N, Aiello D, et al. An insight on type I collagen from horse tendon for the manufacture of implantable devices. Int J Biol Macromol. 2020;154:291–306. doi:10.1016/j.ijbiomac.2020.03.082

70. Voleti PB, Buckley MR, Soslowsky LJ. Tendon healing: Repair and regeneration. Annu Rev Biomed Eng. 2012;14:47–71. doi:10.1146/ANNUREV-BIOENG-071811-150122

71. Buckley MR, Evans EB, Matuszewski PE, et al. Distributions of types I, II and III collagen by region in the human supraspinatus tendon. Connect Tissue Res. 2013;54(6):374–379. doi:10.3109/03008207.2013.847096

72. Smith RK, Birch H, Patterson-Kane J, et al. Should equine athletes commence training during skeletal development?: changes in tendon matrix associated with development, ageing, function and exercise. Equine Vet J Suppl. 1999;30:201–209. doi:10.1111/j.2042-3306.1999.tb05218.x

73. Gonçalves-Neto J, Witzel SS, Teodoro WR, Carvalho-Junior AE, Fernandes TD, Yoshinari HH. Changes in collagen matrix composition in human posterior tibial tendon dysfunction. Joint Bone Spine. 2002;69(2):189–194. doi:10.1016/S1297-319X(02)00369-X

74. Li S, Gong F, Zhou Z, Gong X. Combined Verapamil-Polydopamine Nanoformulation Inhibits Adhesion Formation in Achilles Tendon Injury Using Rat Model. Int J Nanomedicine. 2023;18:115–126. doi:10.2147/IJN.S377600

75. Chien C, Pryce B, Tufa SF, Keene DR, Huang AH. Optimizing a 3D model system for molecular manipulation of tenogenesis. Connect Tissue Res. 2018;59(4):295–308. doi:10.1080/03008207.2017.1383403

76. Atkinson F, Evans R, Guest JE, et al. Cyclical strain improves artificial equine tendon constructs in vitro. J Tissue Eng Regen Med. 2020;14(5):690–700. doi:10.1002/term.3030

77. Schulze-Tanzil G, Mobasheri A, Clegg PD, Sendzik J, John T, Shakibaei M. Cultivation of human tenocytes in high-density culture. Histochem Cell Biol. 2004;122(3):219–228. doi:10.1007/s00418-004-0694-9

78. Petta D, D’Amora U, D’Arrigo D, et al. Musculoskeletal tissues-on-a-chip: Role of natural polymers in reproducing tissue-specific microenvironments. Biofabrication. 2022;14(4). doi:10.1088/1758-5090/ac8767

79. Liu Q, Huang J, Xia J, Liang Y, Li G. Tracking tools of extracellular vesicles for biomedical research. Front Bioeng Biotechnol. 2022;10(November):1–13. doi:10.3389/fbioe.2022.943712

