## Supplementary information for "Allogenic platelet-rich plasma and platelet-rich plasma extracellular vesicles alter the proteome of tenocytes in an in vitro equine model of tendon inflammation: A pilot study"

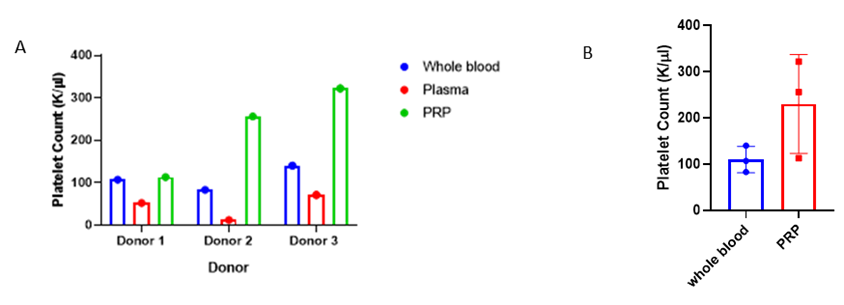

**Figure 1.** (A) The platelet count (K/ul) per anonymised equine donor, from whole blood, plasma and platelet-rich plasma (PRP). (B) The mean (n=3) platelet count in plasma and PRP with standard deviation shown. Both Figure 1A and 1B demonstrate PRP characterisation as previously defined, with pooled PRP (691,000/µl) ) having 2.09 times more platelets when compared to the baseline platelet count from whole blood (330,000/µl).
